# Mechanics of the cellular microenvironment as perceived by cells *in vivo*

**DOI:** 10.1101/2021.01.04.425259

**Authors:** Alessandro Mongera, Marie Pochitaloff, Hannah J. Gustafson, Georgina A. Stooke-Vaughan, Payam Rowghanian, Otger Campàs

## Abstract

Tissue morphogenesis and repair, as well as organ homeostasis, require cells to constantly monitor their 3D microenvironment and adapt their behaviors in response to local biochemical and mechanical cues^1-6^. *In vitro* studies have shown that substrate stiffness and stress relaxation are important mechanical parameters in the control of cell proliferation and differentiation, stem cell maintenance, cell migration ^7-11^, as well as tumor progression and metastasis^12,13^. Yet, the mechanical parameters of the microenvironment that cells perceive *in vivo*, within 3D tissues, remain unknown. In complex materials with strain- and time-dependent material properties, the perceived mechanical parameters depend both on the strain and timescales at which the material is mechanically probed^14^. Here, we quantify *in vivo* and *in situ* the mechanics of the cellular microenvironment that cells probe during vertebrate presomitic mesoderm (PSM) specification. By analyzing the magnitude and dynamics of endogenous, cell-generated strains, we show that individual cells preferentially probe the stiffness associated with deformations of the supracellular, foam-like tissue architecture. We reveal how stress relaxation leads to a perceived microenvironment stiffness that decreases over time, with cells probing the softest regime. While stress relaxation timescales are spatially uniform in the tissue, most mechanical parameters, including those probed by cells, vary along the anteroposterior axis, as mesodermal progenitors commit to different lineages. Understanding the mechanical parameters that cells probe in their native 3D environment is important for quantitative studies of mechanosensation *in vivo*^2-4,6,15^ and can help design scaffolds for tissue engineering applications^16-18^.

---

Cells in tissues constantly make decisions based on the biochemical and mechanical cues they perceive in the local microenvironment^1,3-5^. Using well-controlled mechanical microenvironments in 2D and 3D cell culture systems (e.g., hydrogel scaffolds, purified matrices or substrates with adjustable stiffness), it has been shown that different mechanical inputs affect cell behavior in the absence of instructive biochemical cues^1,11,19^. Notably, stem cell differentiation can be guided by tuning substrate stiffness^10^. Similarly, different threshold values in scaffold elasticity also regulate germ layer specification ^20^, suggesting that in addition to adult tissue homeostasis and regeneration, mechanical cues may be important to control cell fate during embryogenesis^1,3,4,15,21^. Similar to morphogens, spatial gradients of mechanical parameters in embryonic tissues could provide positional information to cells. Moreover, stage-specific changes in tissue mechanics could serve as trigger signals to drive specific cell behaviors, as recently shown *in vivo* for neural crest migration^22^.

The mechanics of the cellular microenvironment has also been shown to be key in bioengineering applications related to stem cell-based tissue regeneration^1,16-18^. In this case, it is important to design synthetic scaffolds with controlled mechanical parameters that mimic the microenvironment that cells perceive *in vivo* during the regenerative response. Most studies have focused on the control of scaffold stiffness, as it has been shown to affect cell differentiation *in vitro*. However, living tissues are often more complex than linear elastic materials, with multiple structures (cytoplasm, cell cortex, adhesion at cell-cell contacts, extracellular matrix, etc.) contributing to their mechanical response at different length- and time-scales. Recent studies using synthetic hydrogel matrices with controlled viscoelastic properties have shown that the timescale of stress relaxation affects cell differentiation, among several other cell behaviors^8,23^. While it is now clear that multiple mechanical parameters influence cell behavior *in vitro*, very little is known about the mechanical parameters of the microenvironment that cells perceive *in vivo*, the structures that cells mechanically probe and how these mechanical cues affect cell behavior within developing 3D tissues. In particular, the mechanical cues that cells experience during embryogenesis, as differentiation of specialized structures takes place, are largely unknown.

During the formation of the vertebrate body axis, mesodermal progenitors located at the posterior end of the body (mesodermal progenitor zone, or MPZ) progressively differentiate into mesodermal cells, establishing the presomitic mesoderm (PSM) and eventually organizing into somites^24^(Fig. 1a). In this process, cells of the posterior paraxial mesoderm modify their transcriptional profile^25^ and progressively acquire an epithelial-like phenotype through a process of mesenchymal-to-epithelial transition (MET) ^26^. Recent *in vivo* mechanical measurements of the (non-linear) tissue mechanical response associated with large tissue deformations (large strains) showed that posterior tissues undergo a transition from a fluid-like state in the MPZ to a solid-like state in the PSM^27^, consistent with their foam-like tissue architecture. In contrast, the (linear) mechanical response of the same tissues to small deformations (small strains) is viscoelastic^28^. These experiments revealed a complex mechanical landscape, with a distinct mechanical response for small vs. large applied strains (plasticity) and a time-dependent response for small tissue deformations (viscoelasticity). Moreover, the associated mechanical parameters (stiffness, viscosity, etc.) in each regime (plastic or viscoelastic) seem to display spatial variations in the tissue^28^. With the mechanical landscape changing with strain, space and time, it is unclear what mechanical parameters cells probe as they differentiate.

**Figure 1.**
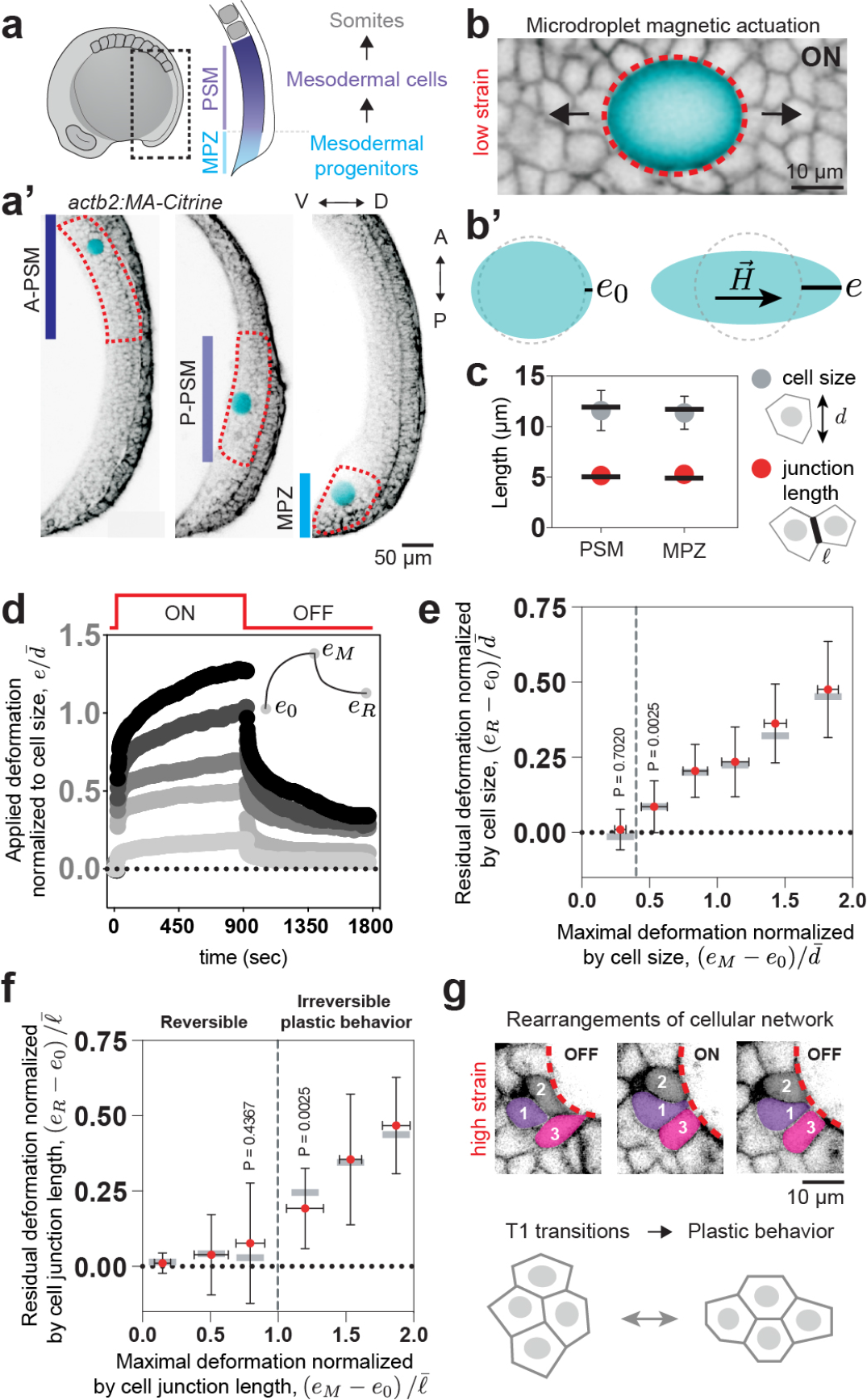
Junctional length establishes the onset of tissue plasticity. **a**, Sketch showing a lateral view of a 10-somite stage embryo highlighting the posterior region of the body (dotted black rectangle) where mesodermal progenitors progressively differentiate as they transit from the mesodermal progenitor zone (MPZ) to the presomitic mesoderm (PSM). **a’**, Confocal sections along the sagittal plane of posterior extending tissues in membrane-labeled *Tg(actb2:MA-Citrine)* embryos (inverted) with inserted ferrofluid droplets (cyan) in different regions along the anteroposterior (AP) axis. The posterior paraxial mesoderm is here divided into three regions (dotted red insets): the anterior presomitic mesoderm (A-PSM), the posterior presomitic mesoderm (P-PSM) and the mesodermal progenitor zone (MPZ). **b**, Confocal section of droplet (cyan) during actuation (magnetic field ON) with small magnetic field leading to small droplet deformation and associated small strains (membrane label; inverted). **b’**, Sketches defining the induced droplet elongation *e* along the direction of the applied magnetic field 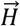 during actuation and the droplet pre-elongation before actuation, *e*_0_. **c**, Mean (dots) and median (line) values of cell size (diameter, *d*; gray) and junctional length 𝓁 (red) in the MPZ and PSM, reanalyzed from Ref. ^27^. **d**, Examples of time evolution of droplet deformation (normalized droplet extension, 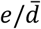, induced strain) during actuation cycles (OFF-ON-OFF; creep experiment) for different values of the applied magnetic field, leading to varying values of maximal droplet elongation *e*_*M*_. The initial and residual droplet deformations, before actuation and after relaxation, *e*_0_ and *e*_*R*_ respectively, are defined in the inset. **e-f**, Residual droplet elongation normalized by average cell size, 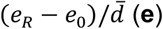, or by average junctional length, 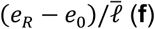, for varying values of the applied maximal droplet elongation (*e*_*M*_ − *e*_0_) normalized by average cell size (**e**) or junctional length (**f**) in the posterior paraxial mesoderm. When the applied maximal droplet elongation exceeds the cell-cell junction length (i.e., for applied strains 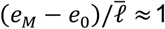; gray dashed line in **f**) the tissue starts to deform irreversibly (plastic deformation). Below this yield strain, the tissue behaves reversely. Circles indicate mean, gray bars median and error bars SD. For cell size normalization N = 105 (divided in the different ranges of applied maximal strain: 7, 14, 37, 22, 16, 9). For junction normalization N = 72 (divided in the different ranges of applied maximal strain: 20, 8, 5, 9, 11, 19). After normalization, data points were grouped in bins of 0.33. Within each bin, the average maximal strain was then calculated. **g**, Snapshots showing confocal sections of tissue next to a droplet (dotted red line) during an actuation cycle (OFF-ON-OFF) with a large applied magnetic field causing large droplet deformations (top). Droplet actuation causes T1 transitions (neighbor exchange) that lead to permanent (irreversible or plastic) changes in the local tissue architecture (bottom).

In order to understand which regime of tissue mechanics cells probe, it is first necessary to characterize the magnitude of deformations (or strains) that mark the transition from a linear viscoelastic response to a plastic regime. To do so, we employed magnetically-responsive microdroplets embedded in the paraxial mesoderm at different locations along the anteroposterior axis^27,28^. By applying a constant, uniform magnetic field to droplets previously inserted between the cells in the tissue (Fig. 1a’; Methods), we induced ellipsoidal droplet deformations characterized by an elongation *e* = *b* − *R* in the direction of the applied magnetic field 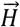 (Fig. 1b,b’) ^28,29^, with *b* and *R* being the droplet semi-axis along the direction of the magnetic field and the radius of the undeformed droplet, respectively. To contextualize droplet deformations in a cellular, foam-like tissue architecture, we normalized the induced droplet elongation, *e*, with the average cell diameter *d* (Fig. 1c; Methods), namely *e*/*d*, which characterizes the applied strain in the material. Increasing magnetic field intensities led to larger droplet deformations (and strains), up to approximately two cell diameters (or 200% strain; Fig. 1d,e). Upon actuation (magnetic field ON; Fig. 1d), droplets progressively elongated from their initial pre-deformed state, characterized by *e*_0_, and reached their maximal elongation *e*_*M*_ just before turning off the magnetic field, at t=15’ (Fig. 1b’,d). After removing the magnetic field (magnetic field OFF; Fig. 1d), capillary stresses pulled the droplet back towards its undeformed, spherical state, progressively reducing the droplet elongation. For values of the applied deformation *e*_*M*_ − *e*_0_ below a threshold value of approximately half the cell size ((*e*_*M*_ − *e*_0_)/*d* ≈ 0.45; Fig. 1e), the process was observed to be largely reversible and the droplet relaxed back to the initial state. In contrast, when the maximal applied deformation *e*_*M*_ − *e*_0_ was above this threshold, the droplet did not fully relax, displaying instead a residual elongation *e*_*R*_ that quantifies the degree of irreversible (plastic) deformation (Fig. 1e). These results show the existence of threshold length scale to induce irreversible changes in the tissue structure and indicate that the length scale controlling the onset of plasticity is smaller than cell size. Normalizing applied deformations with the average cell-cell junction length 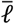 shows that the onset of plastic behavior occurs when applied deformations exceed the cell-cell junction length (Fig. 1c,f), indicating that the characteristic length scale controlling irreversible (plastic) tissue rearrangements is the cell-cell junction length and suggesting that tissue plasticity is due to cell rearrangements (T1 transitions). Indeed, large enough applied deformations induced permanent cell rearrangements in the neighborhood of the droplet (Fig. 1g). This result can be interpreted as the existence of a yield strain (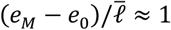; Fig. 1f) in the tissue associated with its foam-like architecture, consistent with previously reported non-linear tissue mechanics^27^.

The mechanical response of the tissue sharply changes when induced deformations are larger than cell-cell contact lengths, indicating that cells will perceive a different mechanical landscape depending on the endogenous strains that they actively generate in the tissue. Since both cell-cell junctions and cellular protrusions can actively generate forces to probe the cellular microenvironment^30-33^ and are mechanosensitive^34,35^, we quantified the strain levels that each of these structures generates. To characterize the strain level generated at cell-cell junctions in the tissue, we monitored the time-evolution of junctional lengths 𝓁 (Fig. 2a) and quantified the endogenous strains by measuring their relative changes over time, namely 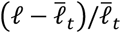, with 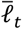 being the time average for individual junctions (Methods). The measured distributions of the magnitude of endogenous junctional strains 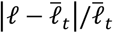 show that progenitor cells in the MPZ generate larger strains than more anterior mesodermal cells of the PSM (Fig. 2b). However, in all cases, the average values of the endogenous strains at cell junctions, which range from 8 to 20% (Fig. 2b), are well below the yield strain (Fig. 1f). In order to quantify the strains generated by protrusions between cells, we performed mosaic labeling of cell membranes and monitored each protrusion length, 𝓁_*p*_, over time (Fig. 2c,d; Methods). Tracking protrusions between cells and calculating the maximal strain levels, 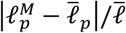 (with 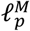 and 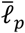 being the maximal and average protrusion lengths; Methods), that protrusions generate shows that protrusion strains are considerably larger in the MPZ than in the PSM (Fig. 2e), with cells generating approximately the same number of protrusions everywhere along the AP axis (Fig. 2f) but with protrusions in the MPZ being consistently longer (Fig. 2g). Yet, all protrusion strains were below the yield strain (Fig. 2e). Altogether, this analysis indicates that both the actomyosin-generated deformations at cell-cell contacts and those generated by protrusions in between cells can only probe the linear mechanical response of the tissue.

**Figure 2.**
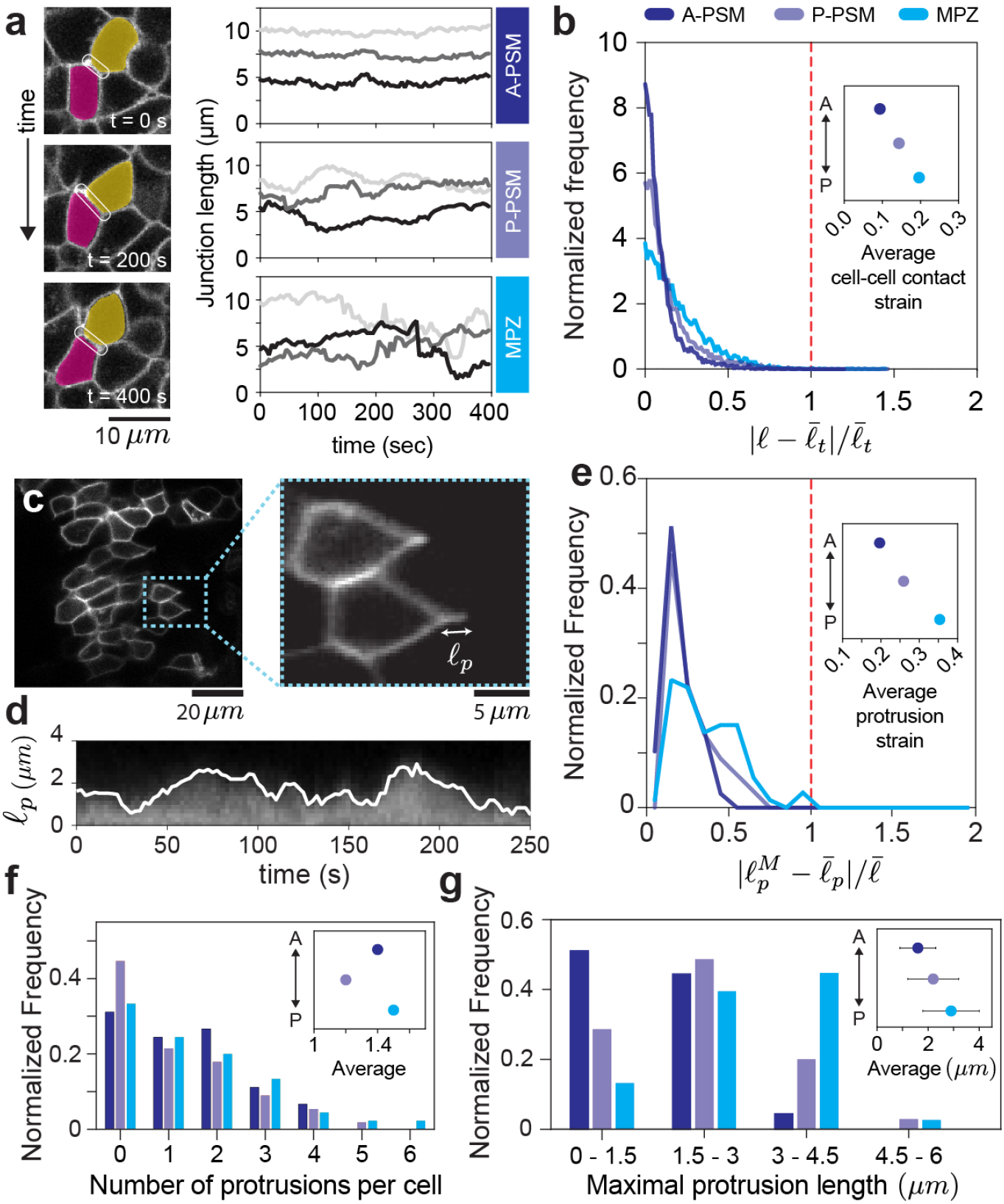
Cells endogenously probe the linear mechanics of the tissue. **a**, Confocal sections showing the temporal changes in cell junction length (white inset) over 400 seconds, and time traces of junction length for cells in different regions of the tissue, showing that junction length is less variable in the A-PSM than in the MPZ. **b**, Normalized frequency (distribution) of the magnitude of relative variations in junction lengths 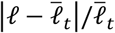 (endogenous applied strain at cell junctions) in the A-PSM and P-PSM (N approximately 3500 junctions in each region; 4 embryos), as well as MPZ (N = 7896; 3 embryos). The average endogenous applied strains at cell junctions (inset) are much smaller than the yield strain in the tissue (red dotted line). **c**, Confocal sections of PSM tissue in mosaic membrane-labeled embryos showing cell protrusions between cells. The length of each protrusion 𝓁_*p*_ (inset) can be measured at each time point. **d**, Kymograph showing the fluorescence intensity along a protrusion path, enabling the determination of the time evolution of the protrusion length (white line), 𝓁_*p*_(*t*). **e**, Normalized frequency of maximal protrusion strains 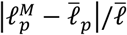 in the different regions. Average protrusion strains (inset) are largest in the MPZ, but much smaller than the yield strain in the tissue (red dotted line). **f-g**, Normalized frequency of the number of protrusions per cell (**f**) and their maximal lengths (**g**), with the average protrusion number per cell and average protrusion length shown in the insets of **f** and **g**, respectively. N = 78, 62, 45 (A-PSM), N = 67, 67, 35 (P-PSM), and N = 73, 66, 38 (MPZ) for **e, f, g**, respectively.

Below the yield strain (linear mechanical response), the tissue material properties are time-dependent^28^ (viscoelastic). It is therefore necessary to know the characteristic timescales at which cells mechanically probe their microenvironment to understand which mechanical parameters they can perceive. Measuring the time evolution of protrusion strain rates, namely (1/𝓁_*p*_) d𝓁_*p*_ /*dt* (Methods), we obtained their distribution in the different regions of the tissue (Fig. 3a,b). The characteristic timescale of protrusions, given by the inverse of the average strain rate, revealed that protrusions in the MPZ were the most dynamic, with characteristic timescales of approximately 1 minute (Fig. 3b, inset). Protrusions in the PSM were slightly slower and displayed characteristic timescales of approximately 2 minutes (Fig. 3b, inset). We then characterized the timescale at which cell-cell junctions probe their microenvironment by measuring their persistence timescale, which we calculated from the autocorrelation function of the junctional length dynamics (Fig. 3a,c; Methods). The average persistence time is of approximately 1 minute in all regions of the tissue (Fig. 3c, inset). To better understand the temporal characteristics of cell-cell junction dynamics, we performed Fourier analysis of the temporal evolution of junctional lengths. Fourier mode amplitudes show that cell-cell junctions preferentially probe the tissue at 1-2 min timescales, irrespective of the location of the cell in the tissue (Fig. 3d), in agreement with the analysis of persistence times of cell-cell junctions (Fig. 3c). Our results indicate that protrusions are most dynamic (∼1 minute timescale) in the MPZ, while cell-cell contact deformations are equally dynamic everywhere in the tissue (∼1 minute persistence time). Either via protrusions or via local junctional contractions, cells preferentially probe their microenvironment at timescales of approximately 1-2 minutes.

**Figure 3.**
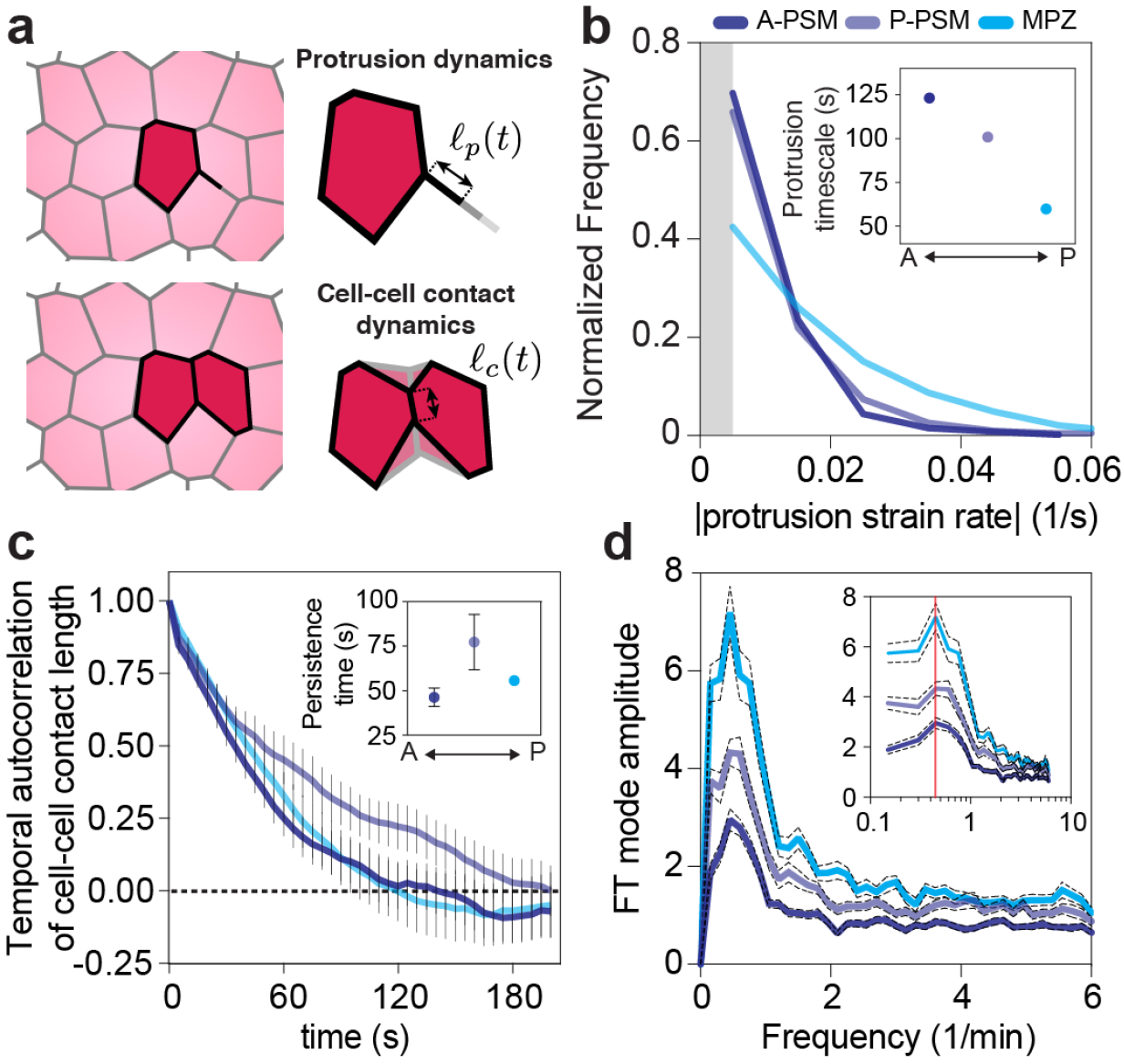
Characteristic timescales of endogenous mechanical probing of the cellular microenvironment. **a**, Sketches of cell protrusions and cell-cell junctions, indicating the measured time-dependent protrusion length and junctional length, 𝓁_*p*_(*t*) and 𝓁(*t*) respectively. **b**, Normalized frequency of the absolute value of the protrusion strain rates, showing minimal protrusion persistence timescales in the MPZ (inset). Limitations in protrusion tracking did not allow measurements of strain rates below approximately 0.005 s^-1^ (gray band). A-PSM (N = 1490), P- PSM (N = 977), and MPZ (N = 391). **c**, Temporal autocorrelation of cell junction length in different regions of the tissue along the AP axis, showing persistence (autocorrelation) timescales of approximately 1 minute (inset). **d**, Fourier Transform (FT) mode amplitudes of the time evolution of junction length in the tissue, for different regions along the AP axis, showing a peak at frequencies of approximately 0.5 min^-1^. N approximately 3500 junctions in A-PSM and P-PSM (4 embryos) and 7896 junctions in MPZ (3 embryos) in **c, d**.

To characterize the mechanics of the cellular microenvironment at strain levels and timescales similar to those endogenously applied by cells (8-35% and 1-2 minutes) in the different regions of the tissue (Fig. 4a), we quantified its time-dependent mechanical response to applied strains ranging between 5-30%. Using magnetically-responsive oil droplets, we performed local creep experiments in the different regions of the paraxial mesoderm, as previously established^28^ (Fig. 1d; Methods). The observed droplet (strain) relaxation dynamics in the studied time period (0-15 min) can be properly accounted for by two distinct relaxation timescales, as previously reported ^28^. To describe the rheological response below yield (no plastic behavior), we used a generalized Maxwell model (Fig. 4b) composed of two Maxwell (viscoelastic) branches, characterized by independent relaxation timescales *τ*_1_ and *τ*_2_, in parallel with an elastic element that accounts for the elastic resistance of the supracellular tissue architecture below yield (before cell rearrangements – plastic events – occur). This is in contrast to previous work^28^, which was unaware of the existence of a yield strain in the tissue, and did not account for the existence of such elastic behavior of the tissue architecture below yield. Measurement of the two stress relaxation timescales shows that these are very different in magnitude, but both uniform along the AP axis (Fig. 4c). The shortest timescale is approximately of 1.6 s, similar to the values previously obtained *in vitro* for the cytoplasm of cells in culture (0.1-1s depending on cell type and conditions^36^) and the stress relaxation timescale of the cytoplasm in the blastomeres of living zebrafish embryos (approximately 2s ^28^). The other stress relaxation timescale is approximately 25s, over 10-fold larger than the shortest (Fig. 4c). This value is close to the previously measured stress relaxation timescale of cellular junctions in epithelial monolayers *in vivo* (∼20s ^37^), the characteristic timescale of actin cortex turnover measured *in vitro* (∼15-45s ^38^) and the timescales of cell-cell adhesion remodeling measured *in vivo* (∼20-40s ^39,40^). These results show that for deformations below the yield strain, stresses in the tissue relax on timescales below 25s approximately.

**Figure 4.**
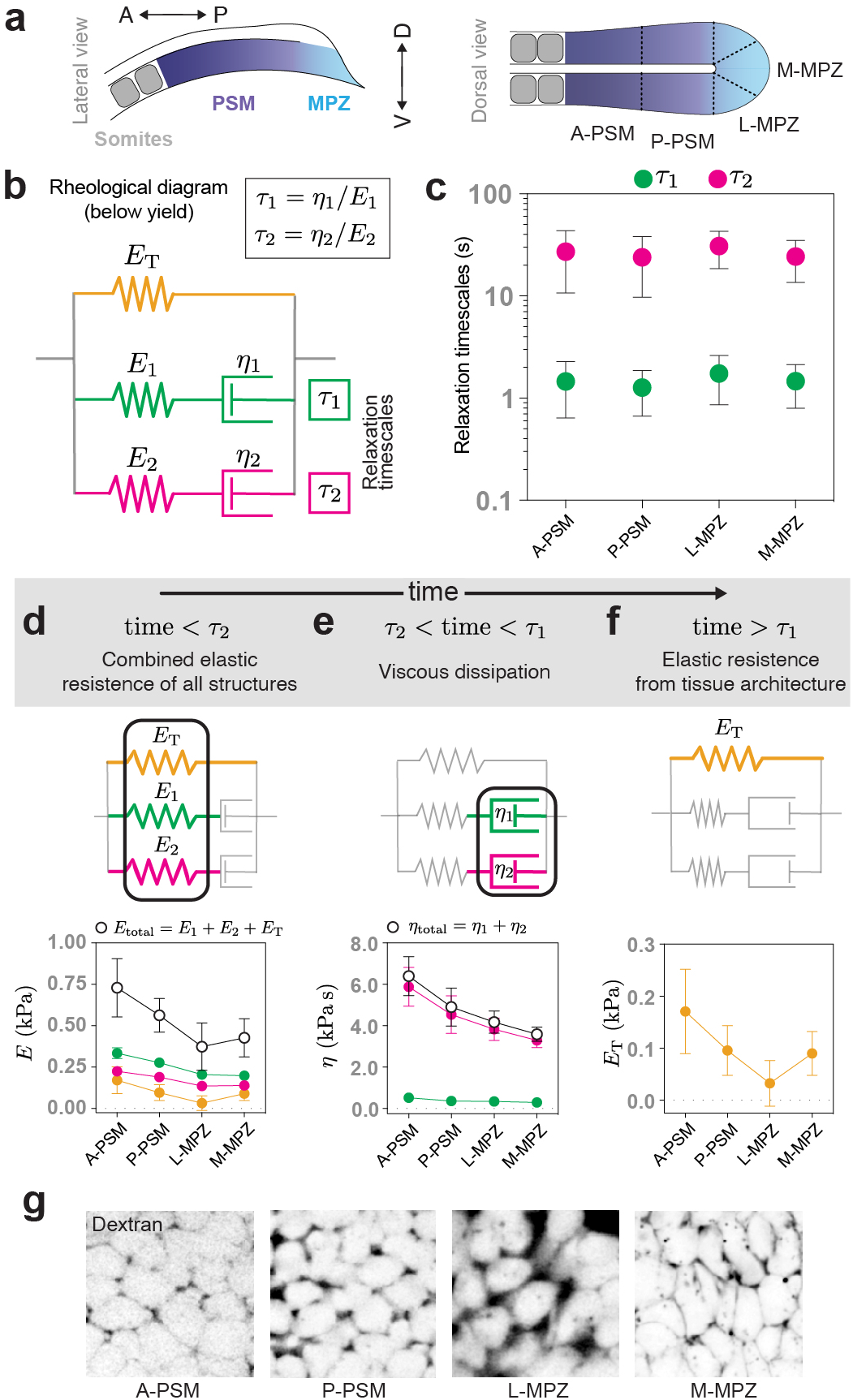
Stress relaxation timescales and time-dependent (linear) mechanical properties of the microenvironment along the AP axis. **a**, Sketch of lateral and dorsal views of the posterior body axis, showing the regions of the tissue in which measurements of linear material properties were performed. The PSM is divided in anterior and posterior parts, A-PSM and P-PSM respectively, and the MPZ is divided in medial and lateral portions, M-MPZ and L-MPZ respectively. **b**, Rheological diagram representing the tissue mechanical response below yield. Two Maxwell branches, characterized by independent stress relaxation timescales *τ*_1_ and *τ*_2_, are in parallel with an elastic element accounting for the stiffness associated with the supracellular foam-like tissue architecture below yield. **c**, Measured values of the two relaxation timescales in the different tissue regions along the AP axis. Mean ± SD. **d-f**, Mechanical properties of the microenvironment at different timescales. All mechanical parameters display a posterior-to-anterior increasing gradient, with the exception of the stiffness of the supracellular tissue architecture, which displays a minimal value in the lateral MPZ. N = 15 for A-PSM, 11 for P-PSM, 17 for L-MPZ, 32 for M-MPZ. Plots show mean ± sem. **g**, Confocal sections revealing the extracellular spaces between cells (fluorescent Dextran; inverted; Methods) in the different regions of the tissue along the AP axis.

To reveal the mechanical parameters that cells probe in the tissue, we measured the different elastic and viscous elements that define the tissue rheological response in the different regions of the tissue (Fig. 4a). Below the smallest stress relaxation timescale (<1s; Fig. 4d), the tissue behaves elastically, with values ranging between 400-800 Pa depending on the tissue region. At intermediate timescales (between ∼1-30s; Fig. 4e), viscous dissipation occurs with two very distinct viscosities. The smallest viscosity, approximately of 80-150 Pa s, is associated with stress relaxation at short timescales (*τ*_1_), whereas the largest one, approximately of 4000 Pa s, is linked to the longer relaxation timescale (*τ*_2_). Finally, above the largest stress relaxation timescale (*t* > *τ*_2_; Fig. 4f) the tissue behaves elastically, with the stiffness values ranging between 30-180 Pa depending on the region of the tissue. While at very long timescales (∼0.5-1h in the MPZ and longer timescales in the PSM) the tissue flows due to plastic cell rearrangements^41^, our results indicate that at timescales of a few minutes (>30s and <10 minutes approximately) the tissue responds to deformations as an elastic material, with its stiffness arising from the resistance to deformations of the foam-like cellular packings that define the tissue architecture. Beyond the time-dependent linear mechanical response, all measured mechanical parameters (stiffnesses and viscosities) monotonically increase away from the posterior end of the body (Fig. 4d-f), with the exception of the stiffness associated with tissue architecture (Fig. 4f), which displays a minimal value in the lateral MPZ (L-MPZ), the region where mesodermal progenitors commit to the mesodermal fate. In foam-like structures, the supracellular stiffness is strongly affected by the amount of extracellular spaces^42^, which are directly related to the physical confinement experienced by cells. Inspection of the extracellular spaces using fluorescent Dextran (Methods) shows that the amount of extracellular spaces is maximal in the L-MPZ (minimal cellular confinement) where the supracellular stiffness displays its smallest value, and minimal in the anterior PSM (A-PSM; maximal cellular confinement) where the supracellular stiffness is the largest (Fig. 2f,g), as expected for foam-like architectures^42^. These results indicate that mesodermal progenitors perceive a minimal stiffness (approximately of 30 Pa) in the L-MPZ (the region with least cellular confinement) as they differentiate into mesodermal cells, which perceive increasing stiffness in the microenvironment during their maturation in the PSM (up to 180 Pa; maximal cellular confinement).

Altogether, these results show that the mechanics of the microenvironment (below yield) displays an effective stiffness that decreases over time, reaching a minimal value for timescales *t* > *τ*_2_ ≈ 25s (Fig. 5). Since cells probe their microenvironment at small strains (<35%; below yield strain) and timescales of approximately 1 minute or above (larger than *τ*_2_) in all regions of the tissue, our results show that cells primarily probe the stiffness associated with the local foam-like tissue architecture (Fig. 5). This suggests that cells may probe their degree of physical confinement, a parameter that has recently been shown to promote somatic-cell reprogramming in 3D environments through an accelerated MET^43^. While our measurements cannot resolve endogenous deformations below our spatial and temporal resolutions (approximately 0.5 μm and 1s; Methods), these small and fast active deformation can only probe the mechanics of smaller subcellular structures, such as the stiffness of the cell cortex. More generally, changes in the stress relaxation timescales of the microenvironment can make cells perceive different stiffnesses at the timescales that they actively probe the microenvironment. This could explain why varying the stress relaxation timescales of the microenvironment can affect cell behavior in similar ways as changing its stiffness. In contrast to the case studied herein, tissues containing substantial extracellular matrix (ECM) will be characterized by additional stress relaxation timescales associated to ECM remodeling^44,45^. To understand what mechanical parameters cells might perceive in ECM-dominated microenvironments, it will be important to know the characteristics of active cellular probing in these contexts and how these compare to the strain- and time-dependent mechanics of the microenvironment^46^.

**Figure 5.**
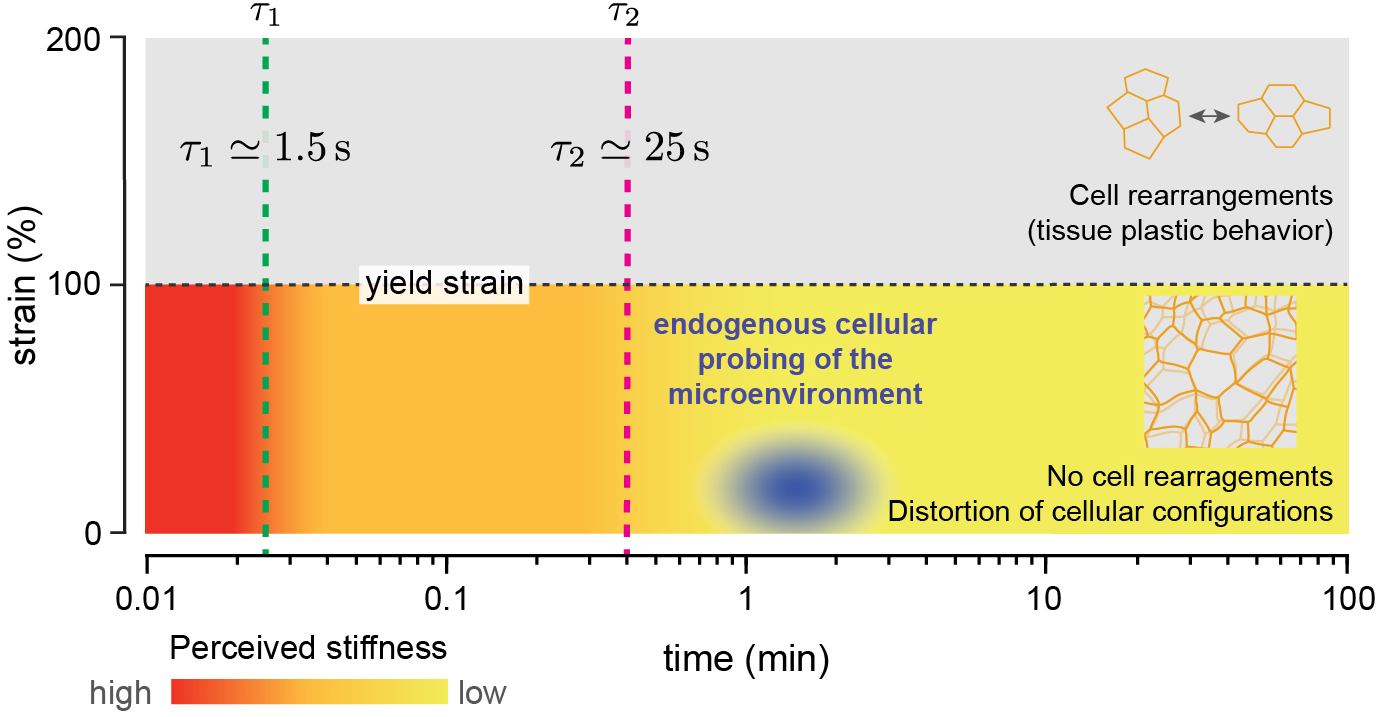
Cells probe the tissue stiffness associated with the local, foam-like architecture of the tissue. Schematic representation of the measured mechanics of the microenvironment at different strains and timescales, showing the different observed regimes and the region of the strain- and time-dependent microenvironment mechanics that is mechanically probed by cells (blue ellipse). Above the yield strain, cell rearrangements (T1 transitions) occur, leading to plastic (irreversible) remodeling of the tissue architecture. Below yield, the tissue maintains its local cellular configurations (no cell rearrangements) and dissipates stresses at different timescales, *τ*_1_ and *τ*_2_ respectively. The perceived microenvironment stiffness below yield (color coded) decreases over time as stresses are dissipated, eventually reaching a constant low value (yellow) associated with the elasticity of deforming cellular packing configurations. Active cellular probing of the microenvironment (blue ellipse) occurs preferentially at timescales (∼1-2 min) longer than all microenvironment relaxation timescales and at small strains (∼10-30%), indicating that cells probe the stiffness associated with the local, foam-like tissue architecture.

Accurate knowledge of what mechanical parameters of the microenvironment cells perceive within living tissues, and how these change in space and time, is essential to understand cellular mechanosensation *in vivo*, during normal development, tissue homeostasis and disease. Moreover, this knowledge can help guide the design of scaffolds for tissue engineering applications that better mimic not only the mechanical parameters that cells perceive *in vivo*, but also the characteristics of the structures responsible for that mechanical response.

## Data Availability Statement

The data that supports these findings are available upon request.

## Acknowledgements

We thank E. Sletten (University of California, Los Angeles) for sharing custom-made fluorinated dyes and S. Megason (Harvard Medical School) for providing *Tg(actb2:MA-Citrine)* embryos. We also thank all past and current members of the Campàs lab for discussions and the CNSI microfluidics facility and the UCSB Animal Research Center for support. PR thanks B. Aigouy (The Developmental Biology Institute of Marseille) for useful technical discussions on Tissue Analyzer. AM was supported by an EMBO Long-Term Fellowship (EMBO ALTF 509-2013), an Errett Discovery Award in Biomedical Research (2015-2016), and an Otis Williams Postdoctoral Fellowship (2016-2017). This work was supported by the National Science Foundation (award number CMMI-1562910), the Eunice Kennedy Shriver National Institute of Child Health and Human Development of the National Institutes of Health (award numbers: R21HD084285 and R01HD095797), and the Deutsche Forschungsgemeinschaft (DFG, German Research Foundation) under Germany’s Excellence Strategy – EXC 2068 – 390729961– Cluster of Excellence Physics of Life of TU Dresden.

## Author Contributions

AM and OC designed research; AM, HJG and GSV performed experiments; AM, HJG, MP and PR analyzed the data; PR and OC performed theoretical interpretation of experiments; AM and OC wrote the paper; OC supervised the project.

## Competing Financial Interests Statement

The authors declare that they have no competing financial interests.

## Online Methods

### Zebrafish husbandry, fish lines, and labeling

Zebrafish (*Danio rerio*) were maintained as previously described^47^. Experiments were performed following all ethical regulations and according to protocols approved by the Institutional Animal Care and Use Committee (IACUC) at the University of California, Santa Barbara. For ubiquitous labeling of cell membranes we used *Tg(actb2:MA-Citrine)* embryos^48^ or embryos injected at 1-cell stage with membrane-GFP mRNA. For mosaic membrane labeling, a small number of cells in wild-type (AB) embryos were injected with 8 – 25 pg of membrane-GFP mRNA at 16 – 32 cell stage.

### Generation and injection of ferrofluid droplets

Ferrofluid droplets were prepared as previously described^28^. Briefly, DFF1 ferrofluid (Ferrotec) was diluted in filtered 3M™ Novec™ 7300 fluorocarbon oil (Ionic Liquid Technologies) at varying concentrations to tune the saturation magnetization of the ferrofluid, thereby allowing variations in droplet deformations (applied strains) for the same applied value of the magnetic field. To prevent non-specific adhesion between cells and droplets, a fluorinated Krytox-PEG(600) surfactant (008-fluorosurfactant, RAN Biotechnologies^49^) was diluted in the ferrofluid at a 2.5% (w/w) concentration. The ferrofluid was calibrated before each experiment as previously described^28^, so that the applied magnetic stresses are known, enabling quantitative experiments. The droplets were directly generated inside the embryos, by micro-injection of the ferrofluid oil in the tissue of interest, as previously described^28^. The required droplet size (of average radius approximately of 20 *μm*) was achieved by modulating the injection pressure and the injection pulse interval. Droplets were injected in the mesodermal progenitor zone (MPZ) at the 4- and 6-somite stages for measurements in the PSM and MPZ, respectively. Experiments were performed at least 2 hours after injection to allow tissue recovery.

### Imaging

Embryos at the 10-somite stage were mounted in 0.8% low-melting agarose and imaged at 25 °C using a laser scanning confocal (LSM 710, Carl Zeiss Inc.). Confocal images of the region of interest in ubiquitous or mosaic membrane-labeled embryos were taken at 2.5s intervals using a 40x water immersion objective (*LD C-Apochromat 1*.*1W, Carl Zeiss Inc*.). Imaging of ferrofluid droplets in the embryo was done as previously described^28^. Ferrofluid droplets were labelled using a custom-synthesized fluorinated Rhodamine dye^50^, which was diluted in the ferrofluid oil at a final concentration of 37 *μM*.

### Magnetic actuation of ferrofluid microdroplets

Actuation of ferrofluid droplets was performed following the previously described protocol^28^. Briefly, a ferrofluid droplet embedded in the tissue was deformed by applying a uniform and constant magnetic field^28^. The magnetic field was applied for 15 minutes, as this timescale is longer than typical cellular processes but minimal tissue rearrangements due to tissue morphogenesis occur within this time period. After 15 minutes the magnetic field was removed and the droplet was monitored during relaxation for an additional 15 minutes, as droplet relaxation occurs over much shorter timescales (Fig. 1d). Upon application of a uniform, constant magnetic field ferrofluid droplets acquire an ellipsoidal shape, elongated along the direction of the applied magnetic field^28^. In our experiment we monitored the time evolution of the droplet deformation (by quantifying the ellipsoidal shape) and measured the initial (*e*_0_), maximal (*e*_*M*_) and residual (*e*_*F*_) droplet elongation in the direction of the applied magnetic field.

### Determination of junctional lengths and their dynamics

To monitor cellular junctions over time, we acquired confocal sections of embryos injected with memGFP mRNA every 5s and for a total period of 30 minutes. The location of cells’ vertices and junctional lengths in the images were detected using Tissue Analyzer^51^. For each embryo, we segmented a region of interest (ROI) along the AP axis for which cell-cell contacts were trackable over a period of at least 400s. We then used the Tissue Analyzer package to obtain the time evolution of contour lengths of cell-cell contacts and the (x,y) positions of the vertices. The average junction length 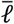 is an ensemble average over different junctions in the tissue.

### Determination of protrusion length and their dynamics

To monitor cell protrusions over time, we acquired time lapses (for 12 min or 18 min at 1 frame every 2.5 s or 16s, respectively) confocal sections of each region of the tissue (MPZ, P-PSM and A-PSM) in wild type embryos with mosaic membrane labeling. Since only a subset of cells were labeled, cell protrusions between cells were visible. Tracking of cell protrusions’ length over time, namely 𝓁_*p*_(*t*), was manually done using ImageJ for each protrusion. The segmented line tool was used to follow each protrusion path, with a large enough width to include the protrusion. The locations of the tip and base (origin of the protrusion on the cell) of each protrusion for each timepoint were determined from the sharp fluorescence changes along the segmented path. The length of the protrusion was then determined for each time point as the difference in length between the tip and base of the protrusion.

### Protrusion strain

To calculate the maximal strains that protrusions can generate, we first determined the maxima in protrusion lengths, namely 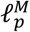, for each protrusion. We excluded maxima separated by less than 0.5 µm or with protrusion length smaller than 0.5 µm because this was close to our spatial resolution. For each protrusion, we then obtained the amplitude of its maximal variations, namely 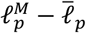, were 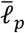 corresponds to the average protrusion length. Since our experimental data showed that the relevant characteristic length scale controlling the onset of plasticity is the junctional length, we obtained the maximal strain applied by protrusions by calculating the ratio of the maximal length variations of protrusions and the average junctional length, namely 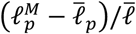. The actual applied strains will, in general, be smaller than the measured values of maximal strains reported here, as the protrusions did not retract instantaneously all their length.

### Protrusion shear rate

Once the time evolution of the protrusion length 𝓁_*p*_(*t*) was determined, we calculated the instantaneous shear rate as (1/𝓁_*p*_) *d*𝓁_*p*_/*dt*. We applied a B-spline using Mathematica (Wolfram) to the measured protrusion length before calculating *d*(ln 𝓁_*p*_(*t*))/*dt* in Matlab (MathWorks). This derivative provides the instantaneous shear rates.

### Persistence timescale of junctional length dynamics

Using the time-series of the contour length of cell junctions (obtained as described in the section *‘Determination of junctional lengths and their dynamics’*), we calculated the temporal autocorrelation function, namely

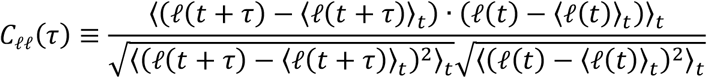

where 𝓁 is the junctional length of any given cell-cell junction, and ⟨𝓁⟩_*t*_ is the time average of a given junctional length. The use of time averages here is because we are interested in the autocorrelation for a given junction. To reduce numerical errors which result from using time averages, these were calculated using time shifted intervals, i.e., each average ⟨·⟩_0_ was evaluated with the data in a time interval (0, *T* − *τ*), with *T* being the duration of the experiment. We analyzed each ROI and obtained the correlation function for each embryo separately. The obtained autocorrelation functions were nearly exponential in all cases. The persistence timescale of the junctional dynamics corresponds to the autocorrelation timescale, which we obtained by fitting an exponential function to the autocorrelation decay. The reported characteristic timescale was obtained from a weighted average of the timescales measured in different embryos, with the weights being the inverse of their variances.

### Fourier transform of junctional length dynamics

We calculate the Fourier transform of the junctional length 𝓁 (obtained as described in the section *‘Determination of junctional lengths and their dynamics’*) using discrete Fast Fourier Transform of the measured junctional length time series. To remove the effect of supracellular, long timescales tissue movements on the junctional lengths, we first filter the junctional length time series using a high-pass filter with a cutoff frequency of approximately 1/6 min^-1^. This suppresses junctional length fluctuations at timescales equal to or larger than 6 minutes. We checked that our results were unchanged by varying the cutoff frequency within a reasonable range. We then subtracted the time average of the filtered signal, namely 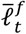, from the time series and calculated the Fast Fourier Transform on the processed signal, namely 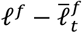.

### Normalized frequency (distribution) of junctional length variations (endogenous strains)

For each one of the time-series of junctional length (obtained as described in the section *‘Determination of junctional lengths and their dynamics’*), we obtain the time average of the junctional length over a period of 400s, namely 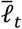, and calculate the relative deviations of the junctional length from its average, namely 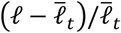. Since actomyosin activity at the cell bjunctions in known to drive the dynamics of cell junctions, these actively generated relative changes in junctional length are a measure of the local, endogenous applied strains. We analyzed 4 ROIs in both A-PSM and P-PSM, and 3 ROIs in the MPZ, all corresponding to different wild type embryos. We obtained the normalized lengths 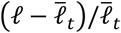 over the time window of a single experiment and combined, for each region, all the values of the strain magnitude (absolute value of 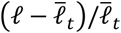 or 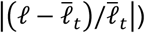, for all times and cell-cell contacts into a single, normalized frequency distribution 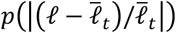 of the endogenous strain magnitude 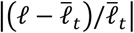.

### Measurement of relaxation time scales and (linear) mechanical properties of the microenvironment

Embryos with a previously injected ferrofluid droplet were mounted for imaging and inspected on the confocal microscope, as described above in ‘*Magnetic actuation of ferrofluid microdroplets*’. Briefly, after the droplet was located, we lowered the magnets array to the distance from the sample that generates the desired magnetic field (and magnetic stress) to create only small droplet deformations leading to applied strains within the 5-20% range, well below the observed yield strain (occurring at 100%). The magnets array was kept at this position for 15 minutes and then moved away from the sample, leading to droplet relaxation. The ferrofluid droplet was imaged for the entire actuation cycle, enabling segmentation of the droplet’s shape in each frame and quantification of deformation dynamics. To obtain all the mechanical parameters in the rheological model (Fig. 2a), we used a previously developed software^28^, which fits the time evolution of the droplet deformation to the mathematical solutions of the rheological model. The mechanical parameters in the rheological model are obtained from the fit parameters, as previously described^28^. Since the droplet capillary stress acts effective as an elastic element acting in parallel of all branches in the generalized Maxwell model^28^, the elastic element associated to the supracellular tissue stiffness (branch 3 in Fig. 2a) and the droplet capillary stress cannot be obtained independently by applying strains below yield. To decouple them, we obtained the effective elastic contribution of the droplet capillary stresses from measurements above yield, as in this case the elastic component of the tissue is not present at long timescales because stresses in the tissue relax via cellular rearrangements. Once the contribution of the capillary stress was known, we removed it from the long timescale elasticity below yield (branch 3 in Fig. 2a). Importantly, the mechanical properties measured with this technique correspond to a local average of the mechanical properties characterizing the surrounding material, along different spatial directions.

### Visualization of extracellular space

To visualize the volume fraction of the extracellular spaces we injected Dextran, Alexa Fluor™ 488 (10000 MW) in the MPZ of 9-somite stage embryos. After 30-45 minutes embryos were mounted and imaged as described above.

### Statistics

In experiments involving zebrafish embryos, the sample size was chosen so that new data points would not change significantly the standard deviation. No samples were excluded from the analysis and the analysis of all the data was done by automated software to ensure full blinding and avoid biases in the analysis. No randomization of the data was used. Mann-Whitney and KS non-parametric tests were used for statistical analysis.

## Notes

### Competing Interest Statement

The authors have declared no competing interest.

